# Phase separation and nucleosome compaction are governed by the same domain of Polycomb Repressive Complex 1

**DOI:** 10.1101/467316

**Authors:** Aaron J. Plys, Christopher P. Davis, Jongmin Kim, Gizem Rizki, Madeline M. Keenen, Sharon K. Marr, Robert E. Kingston

**Affiliations:** Department of Molecular Biology and MGH Research Institute, Massachusetts General Hospital (MGH), Boston, MA, USA; Department of Genetics, Harvard Medical School, Boston, MA, USA; Department of Stem Cell and Regenerative Biology and Harvard Stem Cell Institute, Harvard University, Cambridge, MA, USA; Department of Biochemistry and Biophysics, University of California, San Francisco, San Francisco, CA, USA

## Abstract

Mammalian development requires effective mechanisms to repress genes whose expression would generate inappropriately specified cells. The Polycomb Repressive Complex 1 (PRC1) family complexes are central to maintaining this repression^1^. These include a set of canonical PRC1 complexes that each contain four core proteins, including one from the CBX family. These complexes have previously been shown to reside in membraneless organelles called Polycomb bodies, leading to speculation that canonical PRC1 might be found in a separate phase from the rest of the nucleus^2,3^. We show here that reconstituted PRC1 readily phase separates into droplets *in vitro* at low concentrations and physiological salt conditions. This behavior is driven by the CBX2 subunit. Point mutations in an internal domain of CBX2 eliminate phase separation. These same point mutations eliminate the formation of puncta in cells, and have previously been shown to eliminate nucleosome compaction *in vitro*^4^ and to generate axial patterning defects in mice^5^. Thus, a single domain in CBX2 is required for phase separation and nucleosome compaction, a finding that relates these functions to each other and to proper development.

---

Proper organismal development requires precise regulation of gene expression that is stably maintained. Polycomb-Group (PcG) repressive complexes PRC1 and PRC2 directly act on chromatin to repress key developmental genes and maintain this repressed state throughout development. PRC2 complexes trimethylate lysine 27 on histone H3 (H3K27me3) and this modification recruits canonical PRC1 complexes (Fig. 1a)^1^. Canonical PRC1 complexes contain a CBX protein, which binds to the H3K27me3 mark via a chromodomain at the N-terminus and interacts with RING1b, a key PRC1 protein, via a C-Box at the C-terminus. CBX2, the focus of this study, also contains a positively charged low-complexity disordered region (LCDR). Mutations in this region that reduce the overall positive charge disrupt chromatin compaction *in vitro* and result in axial patterning defects in the mouse^4,5^.

**Figure 1:**
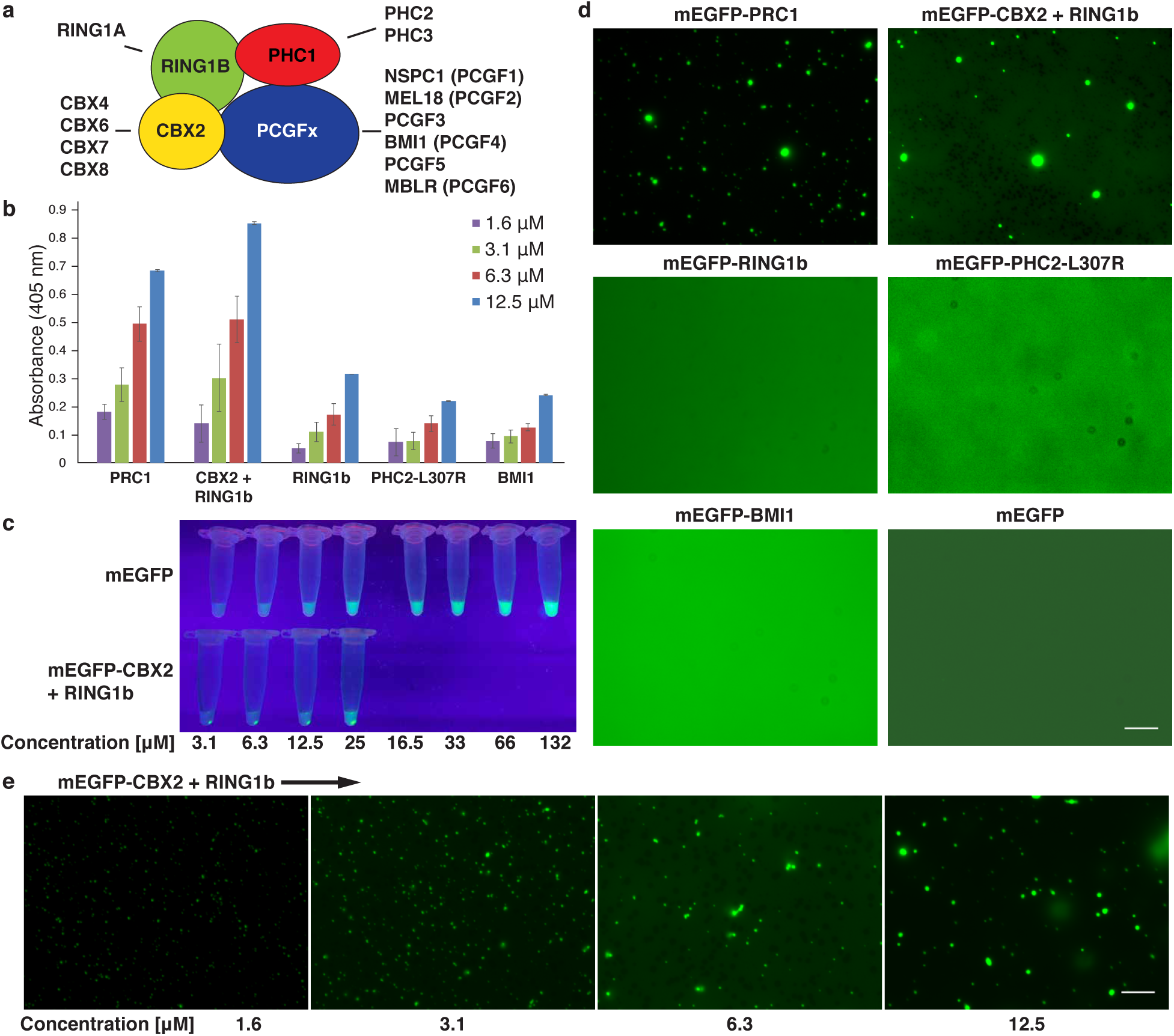
PRC1 phase separates *in vitro*. **a**, Schematic of canonical PRC1 subunits. **b**, Turbid solutions of individual PRC1 subunits at increasing protein concentration were measured at absorbance 405 nm. All proteins are wild type except PHC2-L307R which contains a point mutation in the SAM domain required for expression and purification. **c**, Spin down assay of mEGFP and mEGFP-CBX2 + RING1b to visualize separation of high concentration condensates at increasing protein concentration. **d**, Micrographs of mEGFP-tagged PRC1 subunits at 6.3 μM protein concentration in buffer containing 20 mM HEPES pH 7.9, 100 mM KCl, 1 mM MgSO_4_. For each experiment a representative micrograph from two independent protein preparations is shown. **e**, Micrographs of mEGFP-CBX2 + RING1b at indicated μM protein concentration. Scale bars = 10 μm.

Compaction of chromatin restricts the movement of nucleosomes and makes them less accessible to transcriptional activators. The resulting repressive chromatin state might be further accentuated by compartmentalizing compacted chromatin. Phase separation has been proposed as a mechanism to accomplish this in heterochromatin, based upon the finding that a central component of heterochromatin, HP1a/*α*, phase separates^6,7^. PRC1 is concentrated into nuclear foci called Polycomb bodies^2,3^, one of several classes of ‘membraneless organelles’ that are believed to form by phase separation to enrich and sequester components from bulk solution. Here we show that the CBX2 component of canonical PRC1 can phase separate *in vitro* and generate dynamic puncta in cells. Mutations in CBX2 that impair compaction and proper axial development in mice disrupt phase separation *in vitro* and formation of puncta in cells. This unites, into a single domain within one component of PRC1, the ability to compact nucleosomes and to phase separate, two functions that might coordinate to generate stable repression.

We tested various purified PRC1 protein preparations for turbidity, a known characteristic of phase separated solutions^6,8^ (Fig. 1b). PRC1 formed a turbid solution in a concentration-dependent manner at near-physiological monovalent salt concentration (100 mM KCl). The CBX2-RING1b heterodimer (heterodimerization is necessary to stabilize full length CBX2) displayed turbidity that was more prominent than other individual PRC1 subunits, including RING1b individually. We extended these studies using purified monomeric enhanced GFP (mEGFP)^9^ fusions of PRC1 subunits (Extended Data Fig. 1). After centrifugation of purified protein, mEGFP remained distributed throughout the solution, whereas mEGFP-CBX2 + RING1b coalesced into a protein-rich pellet (Fig. 1c), indicating that mEGFP-CBX2 + RING1b could form a dense phase, separable from bulk solution. Furthermore, fluorescence microscopy revealed the formation of protein-rich foci by purified mEGFP-PRC1 and mEGFP-CBX2 + RING1b, while other PRC1 subunits remained diffusely distributed (Fig. 1d and Extended Data Fig. 2). As seen with other proteins that phase separate^10^, mEGFP-CBX2 + RING1b formed spherical droplets that increase in size as a function of concentration (Fig. 1e). Thus, PRC1 can form phase-separated condensates *in vitro* and CBX2 is a candidate to drive this phase separation.

We examined CBX2 mutants to identify a region needed for phase separation. CBX2 contains a positively charged LCDR (Extended Data Fig. 3a, b), a type of domain often found in proteins that phase separate^11^. This LCDR was previously shown to be critical for the ability of CBX2 to compact nucleosomal arrays *in vitro*^4^ and regulate proper murine development^5^. A paralogous subunit, CBX7, which lacks the ability to compact nucleosomal arrays *in vitro*^4^, does not have a positively charged LCDR. To test the importance of the CBX2 LCDR for phase separation *in vitro*, we purified mEGFP-tagged variants of CBX2 in combination with RING1b that reduce (CBX2-23KRA) or increase (CBX2-DEA) the net positive charge of the region, as well as a heterodimer of mEGFP-CBX7 and RING1b (Fig. 2a and Extended Data Fig. 3c). In contrast to wild-type CBX2, both mEGFP-CBX2-23KRA + RING1b and mEGFP-CBX7 + RING1b failed to form a protein-rich pellet after centrifugation, while mEGFP-CBX2-DEA + RING1b retained the ability to separate from bulk solution (Fig. 2b). In agreement with these data, fluorescence microscopy revealed condensates formed by mEGFP-CBX2 + RING1b and mEGFP-CBX2-DEA + RING1b, whereas mEGFP-CBX2-23KRA + RING1b and mEGFP-CBX7 + RING1b remained diffuse (Fig. 2c and Extended Data Fig. 2). In addition, PRC1 containing mEGFP-CBX2-23KRA showed impaired phase separation relative to PRC1 containing wild-type CBX2, indicating that the LCDR of CBX2 is a driving force for PRC1 phase separation (Fig. 2b, c). Finally, a CBX2 mutation, disrupting only 13 rather than 23 positively charged residues, CBX2-13KRA, also known to impair nucleosome compaction *in vitro* and axial development in mice, failed to phase separate *in vitro* (Extended Data Fig. 4a). We conclude the positive charge within the CBX2 LCDR is critical for phase separation *in vitro*, in addition to its previously described roles in chromatin compaction^4^ and proper axial patterning in mice^5^.

**Figure 2:**
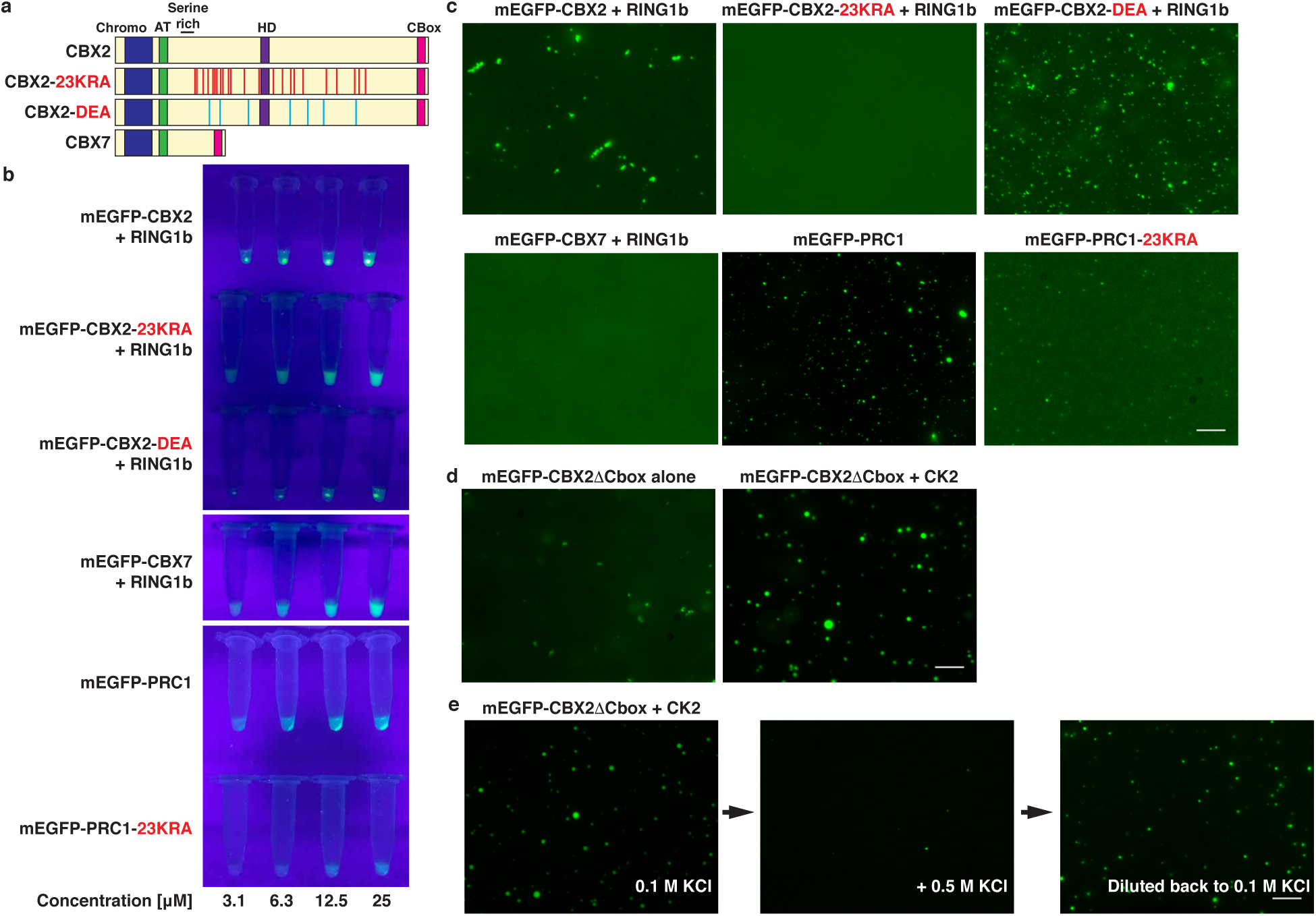
Low-complexity disordered region (LCDR) of CBX2 mediates phase separation *in vitro*. **a**, Schematic of CBX2 mutants and CBX7 protein domains. Point-mutated residues in CBX2-23KRA and CBX2-DEA are highlighted in red and blue, respectively. **b**, Spin down assay of mEGFP-tagged CBX2 mutants and CBX7 + RING1b heterodimers, and full PRC1 complexes to visualize separation of high concentration condensates at increasing protein concentration. **c**, Micrographs of mEGFP-tagged CBX2 mutants and CBX7 + RING1b heterodimers, and full PRC1 complexes all at 6.3 μM. For each experiment a representative micrograph from two independent protein preparations is shown. *d*, Micrographs of mEGFP-CBX2DCbox alone (unphosphorylated) (12.5 μM) or from cells co-expressing catalytic subunits of casein kinase II (CK2) (phosphorylated) (12.5 μM). **e**, Micrographs of salt-dependent reversibility assay. Phosphorylated mEGFP-CBX2ΔCbox (12.5 μM) in buffer containing 100 mM KCl, followed by buffer containing 500 mM KCl and then diluted back to 100 mM KCl. Scale bars = 10 μm.

**Figure 3:**
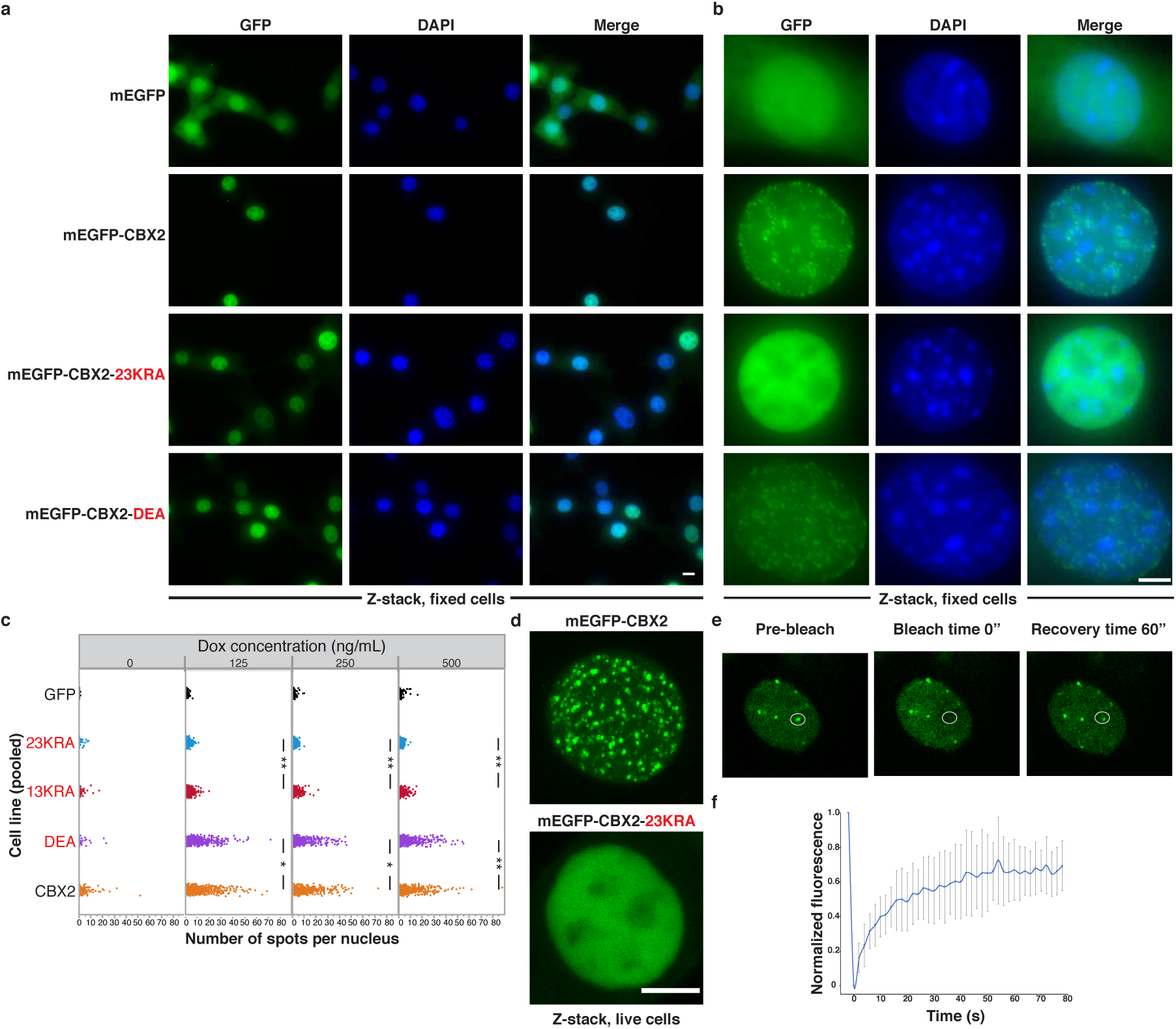
PRC1 proteins form punctate structures *in vivo*. **a**, Representative micrographs of 3T3 fibroblasts after 500 ng/mL doxycycline induction. The mEGFP construct in each cell line is indicated. Column panels from left to right are the GFP channel, DAPI channel and merged images. Scale bar = 10 μm. All cell images are of Z-stacks using formaldehyde fixed cells. **b**, Zoomed in images of representative nuclei from (a) showing puncta or diffuse signal pattern of mEGFP fusion constructs. Scale bar = 5 μm. **c**, High content imaging quantification for the distribution of the number of spots per nucleus from three pooled replicates for indicated 3T3 cell line expressing mEGFP-tagged CBX2 variants at indicated doxycycline concentration. P-value thresholds for statistically significant differences in the distributions of puncta per cell for each doxycycline treatment, as assessed using a two-tailed Mann-Whitney *U* test, are indicated with asterisks (^*^ indicates p-value < 0.01, ^*∗^ indicates p-value ≤ 0.0001). All other combinations (not shown) have a p-value ≤ 0.0001 when doxycycline is present. **d**, Representative micrographs of live 3T3 fibroblasts expressing mEGFP-CBX2 or mEGFP-CBX2-23KRA after 500 ng/mL doxycycline induction. Images are max projections of Z-stacks. Scale bar = 5 μm. **e**, Representative images of FRAP experiment with mEGFP-CBX2 after 500 ng/mL doxycycline induction in 3T3 fibroblasts. White circle indicates bleached region of interest (ROI) over puncta. **f**, Quantification of FRAP data of mEGFP-CBX2. FRAP curve was generated as the mean of n = 15 puncta. Error bars represent standard deviation.

The observation that mutations in positively charged residues disrupt phase separation raised the hypothesis that negatively charged residues in CBX2 might form multivalent interactions with the positive residues. Mutation of negative residues within the LCDR (CBX2-DEA) did not impact phase separation, leading us to consider other sources of negative charge. Phosphorylation increases negative charge and modulates phase separation of proteins both positively and negatively^6,7,12–15^. Serine residues in CBX2 are phosphorylated *in vivo* and targeted by casein kinase II (CK2) *in vitro*^16^. We tested a role for phosphorylation in condensate formation by using *E.coli* to express a truncated, non-phosphorylated, form of CBX2 (mEGFP-CBX2DCbox) stable in the absence of RING1b. We also co-expressed this protein with the catalytic subunits of CK2 to generate phosphorylated mEGFP-CBX2DCbox. Phosphorylation was validated by mass spectrometry (Extended Data Fig. 5, Supplementary Table 1). Phosphorylated mEGFP-CBX2DCbox formed spherical droplets, distinct in size and shape from the more diffuse signal and small non-spherical aggregates formed by unphosphorylated mEGFP-CBX2ΔCbox (Fig. 2d). Thus, phosphorylation of CBX2 increases the ability of CBX2 to phase separate, suggesting a role for electrostatic interactions within CBX2 in driving condensate formation.

To determine whether the droplets formed by mEGFP-CBX2ΔCbox were solid aggregates or reversible liquid condensates, we performed a salt-dependent reversibility assay. Droplets were formed and visualized as described above (Fig. 2d, e) and the salt concentration was then increased to 500 mM KCl. At higher salt, the preformed droplets drastically reduced in number and size (Fig. 2e). Reducing the salt concentration to 100 mM KCl resulted in reformation of droplets, albeit smaller due to reduced protein concentration (Fig. 2e). These results support the hypothesis that reversible electrostatic interactions between phosphorylated serines and positively charged residues are necessary for phase separation of CBX2 *in vitro*.

As mutations in the CBX2 LCDR impair its ability to phase separate *in vitro*, we assessed the impact of these mutations on the morphology of structures formed by PRC1 *in vivo.* We expressed different mEGFP-CBX2 variants under a doxycycline-inducible promoter in 3T3 fibroblasts. Induction of mEGFP expression produced diffuse signal throughout the nucleus and cytoplasm (Fig. 3a, b). In contrast, mEGFP-CBX2 formed nuclear puncta, similar to those previously observed for PRC1 in other cell types^2,3,17,18^. mEGFP-CBX2-KRA mutants failed to form nuclear puncta, while the mEGFP-CBX2-DEA mutant formed puncta similar to those seen for wild-type CBX2 (Fig 3a, b and Extended Data Fig. 4b, c). Quantification of puncta in mEGFP-CBX2 and mEGFP-CBX2-23KRA expressing nuclei revealed a clear difference in total number and distribution across a range of doxycycline concentrations (Fig. 3c, Extended Data Fig. 6). There was also a significantly higher number of puncta in mEGFP-CBX2-13KRA compared to mEGFP-CBX2-23KRA expressing nuclei. This intermediate defect for CBX2-13KRA mirrors the less severe defects in chromatin compaction activity *in vitro* and *in vivo* axial patterning phenotype for this mutant relative to CBX2-23KRA. To address whether these puncta contain canonical PRC1 subunits, we used co-immunoprecipitation. This showed that both mEGFP-CBX2 and mEGFP-CBX2-23KRA interacted with other PRC1 subunits *in vivo* (Extended Data Fig. 7a, Supplementary Table 2). Co-immunofluorescence of RING1b revealed extensive co-localization with mEGFP-CBX2 (Extended Data Fig. 7b) in 3T3 fibroblasts, which do not endogenously express CBX2 (Extended Data Fig. 7c). Thus, the puncta visualized by mEGFP-CBX2 contained PRC1. These *in vivo* results recapitulate the findings of our *in vitro* assays and underscore the importance of positively charged residues in the CBX2 LCDR for PRC1 phase separation.

Phase-separated condensates undergoing demixing with the surrounding aqueous environment display a rapid exchange of interacting components^19^. To interrogate the dynamics of nuclear puncta formed by CBX2 *in vivo*, we performed live cell microscopy of 3T3 fibroblasts expressing mEGFP-CBX2 and mEGFP-CBX2-23KRA (Figure 3d and Extended Data Fig. 8). As seen in formaldehyde-fixed cells, mEGFP-CBX2 organized into puncta whereas mEGFP-CBX2-23KRA remained diffusely distributed throughout the nucleus. To examine whether mEGFP-CBX2 puncta behave as liquid-like condensates, we performed fluorescence recovery after photobleaching (FRAP). Upon photobleaching, mEGFP-CBX2 puncta rapidly recover fluorescence within 60 seconds (Fig. 3e, f). Consistent with these nuclear puncta behaving as phase separated condensates, we observed a rapid loss of puncta upon addition of 1,6-hexanediol, as observed for other phase separated bodies (Extended Data Fig. 9)^7,12,20–22^. We conclude that CBX2 within puncta can readily exchange with free CBX2 in the surrounding environment, consistent with the properties of a liquid-like condensate.

Phase separation can facilitate inclusion or exclusion of macromolecules from the protein-dense phase, creating a mechanism to compartmentalize biochemical activities^11^. We tested whether ligands of PRC1, including DNA, RNA, and nucleosomal arrays, could incorporate into PRC1 condensates *in vitro.* We generated polynucleosomal templates using Cy5-labeled G5E4 DNA^23^, either with heterogeneously modified polynucleosomes or with polynucleosomes containing an H3K27me3 analog^24^. We also included Cy5-labeled G5E4 DNA alone, as well as Cy5-labeled CAT7 RNA previously shown to associate with PRC1^25^. We monitored incorporation of these ligands into PRC1 condensates using fluorescence microscopy. All four ligands were incorporated into condensates formed by mEGFP-CBX2 + RING1b (Fig. 4a) and mEGFP-PRC1 (Fig. 4b), whereas free Cy5 dye was not found within the condensate phase. Ligands incorporated into PRC1 condensates regardless of whether they were added to preformed droplets (Extended Data Fig. 10) or included during droplet formation (Fig. 4a). Thus, PRC1 condensates partition with physiologically relevant ligands, suggesting a mechanism to compartmentalize these interactions *in vivo*.

**Figure 4:**
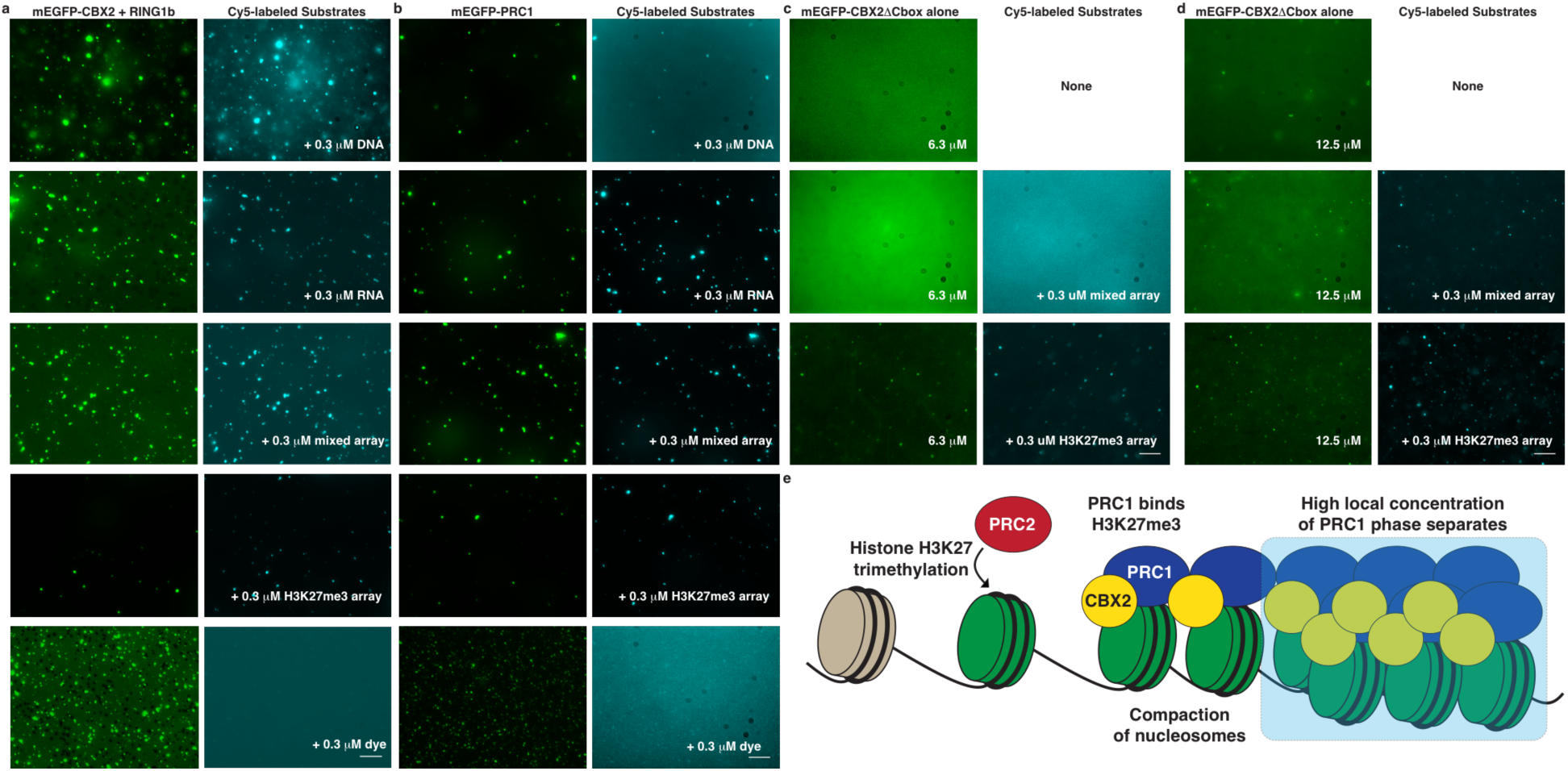
PRC1 ligands partition into condensates with PRC1. **a**, Micrographs of mEGFP-CBX2 + RING1b (6.3 μM) with indicated Cy5-labeled substrate (0.3 μM). Left panels are GFP channel and right panels are Cy5 channel. **b**, Micrographs of mEGFP-PRC1 (6.3 μM) with indicated Cy5-labeled substrates (0.3 μM). Panels are the same as for (A). **c**, Micrographs of unphosphorylated mEGFP-CBX2DCbox (6.3 μM) alone (top) or with indicated Cy5-labeled heterogeneously modified (mixed) or H3K27me3-modified polynucleosomes (0.3 μM). **d**, Same as in (c) with higher concentration of unphosphorylated mEGFP-CBX2DCbox (12.5 μM). Scale bar = 10 μm. **e**, Model of PRC1 nucleosome compaction and phase separation.

The bacterially produced unphosphorylated mEGFP-CBX2DCbox did not phase separate by itself (Fig. 2d) but can compact nucleosomal templates^4^, indicating a possible difference between these activities. We tested the ability of this protein to phase separate under conditions where compaction can occur, which requires the presence of nucleosomal arrays. Nucleosomal arrays might increase the effective local concentration of this protein, and thus might enhance interactions required for phase separation. Incubating unphosphorylated mEGFP-CBX2DCbox with nucleosomal arrays resulted in condensate formation (Fig. 4c, d). Furthermore, we observed that nucleosome arrays containing an H3K27me3 analog, which bind with higher affinity to the CBX2 protein^26^, were more proficient at inducing condensate formation at lower protein concentration. This result is consistent with the hypothesis that phase separation requires a high local concentration of CBX2 protein that can be driven by phosphorylation to increase electrostatic interactions, or to a lesser extent by the addition of nucleosomal arrays to provide a scaffold to facilitate CBX2 interactions.

We show that the abilities of PRC1 to phase separate and to compact nucleosomes both require the LCDR of CBX2 and are inhibited by mutation of basic residues. As this domain lies downstream of the chromodomain that binds H3K27me3, several Polycomb-Group functions are combined into a single protein. A simple hypothesis is that nucleosome compaction and phase separation are manifestations of the same phenomenon, and that compacted and phase separated H3K27me3 nucleosomes are separated from the rest of the nucleus. It has previously been shown that the Polyhomeotic (PH) subunit of PRC1 mediates subnuclear clustering through polymerization of its SAM domain^27–29^. We cannot rule out the possibility that PHC1/2 contribute to PRC1 phase separation as only mutants defective in polymerization were tested here for technical reasons. Notably, PHC1 contains an LCDR rich in glutamine residues, which are highly represented in the LCDRs of other proteins that phase separate^30^. Altogether, we propose a model whereby PRC1 compacts nucleosomes and organizes them into phase separated subnuclear condensates in a concerted manner to efficiently and stably repress transcription (Fig. 4e). This raises questions concerning the state of phase separation by PRC1 during replication and cell division and whether phase separation plays a role in the stable inheritance of repression during differentiation.

## Methods

### Cell Culture

NIH-3T3 fibroblasts (ATCC) were cultured in DMEM supplemented with fetal calf serum to 10% concentration (v/v) and 25 mM HEPES pH 7.5. HEK293T (ATCC) cells were cultured in IMDM supplemented with fetal bovine serum to 10% concentration (v/v). CJ7 (a gift of Stuart Orkin^31^) mouse embryonic stem cells (mESCs) were cultured on a layer of mitotically-inactivated PMEF-N mouse embryonic fibroblasts (Millipore) in DMEM supplemented with fetal bovine serum (Hyclone) to 15% concentration (v/v), 1X L-glutamine, 1X penicillin/streptomycin, and 10 ng/mL leukemia inhibitory factor (LIF). CJ7 media was exchanged daily. NIH-3T3, HEK293T, and CJ7 cells were maintained in a humidified incubator at 37°C with 5% CO_2_. Sf9 cells were maintained in either Hyclone CCM3 or ESF 921 (Expression Systems) media at 27°C in a shaking incubator.

### Isolation of primary tissue from mice

All animal procedures were performed according to NIH guidelines and approved by the Committee on Animal Care at Massachusetts General Hospital and Harvard University. Embryonic day 11.5 (E11.5) mouse embryos were isolated from crosses between C57BL/6 mice heterozygous for a deletion in *Cbx2*, producing *Cbx2*^+/+^, *Cbx2*^+/-^, and *Cbx2*^−/−^ progeny. The *Cbx2* deletion mutant mouse lines arose from CRISPR-mediated modification of *Cbx2* without homology repair during generation of *Cbx2*-*KRA* mice^5^, resulting in a premature stop codon at amino acid position 171 (missense after amino acid 169).

### Expression and purification of proteins from Sf9 cells

For expression of individual PRC1 subunits from Sf9 cells, cDNAs encoding various PRC1 subunits were cloned into pFastbac1, incorporating an N-terminal FLAG tag. For expression of monomeric enhanced GFP (mEGFP) and individual mEGFP-tagged PRC1 subunits from Sf9 cells, cDNAs encoding various PRC1 subunits (excluded for mEGFP alone) were cloned into pFastbac1, incorporating a FLAG tag, the cDNA encoding mEGFP (Addgene plasmid 18696; a gift of Karel Svoboda), and a seven amino acid linker (GSAAAGS) at the N-terminus. These constructs were used to generate baculovirus using the Bac-to-Bac system (Thermo Fisher Scientific). Sf9 cells were infected with baculovirus and incubated with shaking for 72 hours at 27°C to express proteins. For expression of full PRC1 complex, only the CBX2 subunit was FLAG-tagged. Sf9 cells were harvested by centrifugation and used to prepare nuclear extracts as previously described^32^. Nuclear extract was incubated with anti-FLAG M2 affinity resin (Sigma) for 2 hours and then washed with BC buffer (20 mM HEPES at pH 7.9, 0.2 mM EDTA, 20% glycerol, 0.05% NP-40, 0.5 mM DTT, 0.1 mM PMSF, cOmplete EDTA-free protease inhibitor (Roche)) containing 300 mM KCl. Resin was washed with BC buffer containing increasing concentrations (300-600-1200-2000 mM) of KCl, and then washed with BC buffer in descending order of KCl concentration to 300 mM KCl. Proteins were eluted from resin using BC buffer containing 300 mM KCl and 0.8 mg/mL FLAG peptide. Purified protein was concentrated using Amicon Ultra-4 centrifugal filter units and quantified by Bradford assay. The purity of complexes was assessed by Coomassie staining.

### Expression and purification of proteins from *E. coli*

For expression of mEGFP-CBX2ΔCbox from *E. coli* cells, cDNA encoding CBX2ΔCbox was cloned into pET15b, incorporating a FLAG tag and the cDNA encoding mEGFP at the N-terminus. This vector was used to transform Rosetta (DE3) pLysS *E. coli* for protein purification. Phosphorylated mEGFP-CBX2ΔCbox was obtained by co-expression with the catalytic subunits of CKII in a pRSF-Duet vector. Cells were grown to an OD 0.6 at 37°C in 2-YT with 50 μg/mL carbenicillin and 25 μg/mL chloramphenicol. For co-expression with pRSF-Duet CKII vector, 25 μg/mL kanamycin was added. Cells were induced with 0.5 mM isopropyl β-D-1-thiogalactopyranoside overnight at 18°C. Cell extracts were prepared as previously described^4^. Briefly, harvested cells were resuspended in lysis buffer (50 mM HEPES at pH 7.5, 0.5 mM EDTA, 1.6 M KCl, 20% glycerol, 0.5 mM MgCl_2_, 0.05% NP-40, 1 mg/mL lysozyme, 1 mM DTT, protease inhibitors). The cells were taken through three freeze–thaw cycles, then sonicated to shear DNA before centrifugation at 25,000g for 20 minutes to remove debris. Five percent polyethelenimine (PEI) in 20 mM HEPES pH 7.5 was added dropwise to the supernatant while stirring to a final concentration of 0.15%, and stirred an additional 30 minutes. The precipitated nucleic acid was removed by centrifugation at 25,000g for 20 minutes. Extracts were bound to M2 resin and protein purification was carried out as described for Sf9 cells.

### Turbidity assay

To measure turbidity of purified proteins, concentrated proteins were serially diluted to the specified concentrations into buffer containing a final concentration of 20 mM HEPES pH 7.9, 100 mM KCl, and 1 mM MgSO_4_. Diluted proteins were loaded into a clear bottom 384-well plate (Corning), and absorbance at 405 nm was measured using a Spectramax M3 plate reader. Turbidity measurements reflect the average of 3 samples.

### Centrifugation assay

Serial dilutions of mEGFP-tagged proteins were performed in 0.5 mL microcentrifuge tubes as described above for untagged proteins in the turbidity assay. The samples were incubated at room temperature for 5 minutes and then centrifuged at 10,000g for 5 minutes. Material was visualized under UV light.

### Fluorescence microscopy of *in vitro* protein condensates

Prior to imaging, purified mEGFP-tagged proteins were diluted to specified concentrations into buffer containing a final concentration of 20 mM HEPES pH 7.9, 100 mM KCl, and 1 mM MgSO_4_ and spotted on glass slides with coverslips. Proteins were imaged with a Nikon 90i Eclipse epifluorescence microscope equipped with an Orca ER camera (Hamamatsu) using a 100X oil objective and Volocity software (Perkin Elmer). Images in figures were prepared using Fiji software.

### Generation of cell lines for doxycycline-inducible expression of mEGFP-CBX2 variants

cDNAs encoding mEGFP and mEGFP-CBX2 variants were cloned into a modified pTRIPZ vector (Dharmacon). In the modified vector, the RFP and shRNA encoding segments were removed by restriction digest with AgeI and MluI and replaced with a multiple cloning site. pTRIPZ vectors expressing mEGFP and mEGFP-CBX2 variants were transfected into HEK293T in combination with pCMV-dR8.91 containing gag, pol, and rev genes and pMD2.G encoding VSV-G envelope protein using TransIT-Lenti transfection reagent (Mirus). After 48 hours, medium was collected and filtered through a 0.45 μm filter. Filtered medium was concentrated using Lenti-X concentrator (Takara) and concentrated lentivirus was resuspended in Opti-MEM (Thermo Fisher Scientific). NIH-3T3 fibroblasts were transduced with lentivirus at low multiplicity of infection. After 48 hours, transduced cells were selected with puromycin at a final concentration of 2 μg/mL. After selection, stably transduced 3T3 cells were maintained as detailed above.

### Fluorescence microscopy of doxycycline-inducible cell lines

For fixed cell experiments, transduced 3T3 fibroblasts were grown on coverslips. To induce expression of mEGFP and mEGFP-CBX2 variant fusions, media containing the indicated concentration of doxycycline (Sigma) was added for 24 hours. Coverslips were washed with PBS and then crosslinked with 4% formaldehyde in PBS for 15 minutes. The formaldehyde was removed and coverslips were washed twice with PBS. Coverslips were mounted on slides with mounting media containing DAPI (Vector Laboratories, Vectashield H-1200) and imaged with a Nikon 90i Eclipse microscope equipped with an Orca ER camera (Hamamatsu) using a 60X oil objective and Volocity software (Perkin Elmer). A Z-stack of images was collected with 0.2 μm spacing and collapsed using maximum intensity. Images in figures were prepared using Fiji software. For live cell imaging, cells were grown on 35 mm glass bottom fluorodishes (WPI) in phenol red free media and induced with 500 ng/mL of doxycycline for 24 hours. Cells were imaged using a Nikon A1R laser-scanning confocal inverted microscope equipped with a thermostatically controlled stage maintained at 37°C with a 63X oil immersion objective. A Z-stack of images was collected with 0.5 μm spacing and collapsed using maximum intensity. For high content imaging and unbiased quantification of nuclear puncta, transduced 3T3 fibroblasts were grown in black-walled, poly-L-lysine-coated 96 well microplates (Greiner, 655090) and induced with indicated concentration of doxycycline for 24 hours. The cells were fixed as described above for coverslips and stained with Hoechst 33342 (Thermo Fisher Scientific, H3570). Images were acquired on the Opera Phenix High Content Screening System (Perkin Elmer). Confocal images with 4 stacks per field and 28 fields per well were automatically acquired using a 63X water objective. Three replicates per cell line and doxycycline concentration were included in each experiment. At least 500 cells were analyzed for each experimental group. Image segmentation, nuclei and spot identification per cell, and quantification was performed using the Columbus Data Storage and Analysis System (Perkin Elmer). After running the spot-finding script on wells without doxycycline, the raw spot intensities were averaged and standard deviation calculated. Mean of intensities plus two standard deviations was applied as the intensity threshold for identifying positive spots in all the wells. Statistically significant differences in the distributions of puncta per cell for each doxycycline treatment were assessed using a two-tailed Mann-Whitney *U* test.

### Co-immunoprecipitation of mEGFP-CBX2 variants

Transduced 3T3 fibroblasts containing mEGFP, mEGFP-CBX2 or mEGFP-CBX2-23KRA were grown to 80% confluency in 15 cm tissue culture dishes. Media containing 500 ng/mL of doxycycline was added for 24 hours. Cells were washed with PBS and collected using a cell scraper. Nuclear extracts were prepared as previously described^32^. Protein levels in nuclear extracts were measured on a Nanodrop using A280. Equal protein mass between samples was used in subsequent co-immunoprecipitation (co-IP). 1% volume was saved as input. For co-IP, magnetic protein A beads (Invitrogen) were pre-equilibrated in BC buffer containing 300 mM KCl and 0.05% NP-40. Washes were performed on a magnetic rack. 2.5 μg of GFP antisera (Abcam, ab290) for each IP was conjugated to pre-equilibrated beads by incubating for 1 hour at 4°C. GFP antisera conjugated beads were washed three times with BC buffer containing 300 mM KCl and 0.05% NP-40 and mixed with nuclear extracts for 2 hours at 4°C. IPs were washed three times with BC buffer containing 300 mM KCl and 0.05% NP-40 and resuspended in 1X SDS-sample buffer. Samples were heated to 95°C for 5 minutes and supernatant was loaded onto an SDS 4-20% polyacrylamide gel (Biorad). Samples were either processed for mass spectrometry or immunoblotting.

### Mass spectrometry

To detect proteins associated with mEGFP-CBX2 variants by mass spectrometry, co-immunoprecipitated material was run on an SDS polyacrylamide gel and Coomassie stained. Four gel sections were excised for each immunoprecipitation. Gel sections were minced and subjected to a modified in-gel trypsin digestion procedure^33^. Gel pieces were dehydrated with acetonitrile and dried to completion in a SpeedVac. Gel pieces were rehydrated with 50 mM ammonium bicarbonate supplemented with 12.5 ng/μl modified sequencing-grade trypsin (Promega) at 4°C. Rehydrated samples were then incubated at 37°C overnight. Peptides were extracted by removing the ammonium bicarbonate solution, washed with a solution of 50% acetonitrile and 1% formic acid, and dried in a SpeedVac. Dried samples were reconstituted in HPLC solvent A (2.5% acetonitrile, 0.1% formic acid). Samples were loaded onto a reverse-phase HPLC capillary column packed with 2.6 μm C18 spherical silica beads into a fused silica capillary^34^. After gradient formation, peptides were eluted with increasing concentrations of HPLC solvent B (97.5% acetonitrile, 0.1% formic acid). Eluted peptides were subjected to electrospray ionization and entered an LTQ Orbitrap Velos Pro ion-trap mass spectrometer (Thermo Fisher Scientific). Peptides were detected, isolated, and fragmented to produce a tandem mass spectrum of specific fragment ions for each peptide. Peptide sequences were identified using Sequest^35^. All databases include a reversed version of all the sequences. Data were filtered to between a 1-2% peptide false discovery rate.

To identify phosphorylated residues within CBX2 purified from *E. coli* and Sf9 cells, purified protein was run on an SDS polyacrylamide gel and Coomassie stained. A band corresponding to the molecular weight of tagged CBX2 was excised from the gel and analyzed by mass spectrometry as above, with the following alterations. Prior to in-gel trypsin digestion, minced gel pieces were reduced with 1 mM DTT for 30 minutes at 60°C followed by alkylation with 5 mM iodoacetamide for 15 minutes in the dark at room temperature. During mass spectrometry analysis, a modification of 79.9663 mass units to serine, threonine, and tyrosine was included in the database searches to determine phosphopeptides. Phosphorylation assignments were determined by the Ascore algorithm^36^.

### Immunofluorescence

Fixed cells on coverslips were washed once with PBS, then permeabilized in PBS with 0.2% Triton X-100 for 15 minutes. After permeabilization, cells on the coverslip were blocked for 30 minutes with incubation solution (PBS with 3% BSA, 0.05% Triton X-100). Coverslips were then incubated with incubation solution containing primary antibodies (anti-RING1b (Bethyl, A302-869A, 1:4500) or anti-RING1b (Abcam, ab3832, 1:900)) overnight at 4°C in the dark to minimize bleaching of GFP fluorescence. Coverslips were washed three times with PBS with 0.1% Tween-20, then incubated with incubation solution containing secondary antibodies (Alexa-568 conjugated anti-rabbit, or Cy3 conjugated anti-goat, both 1:500) for two hours in the dark. After three washes with PBS containing 0.1% Tween-20, coverslips were rinsed with distilled water and mounted on slides with mounting medium containing DAPI. All incubations and washes were done at room temperature except incubation with primary antibody. Slides were imaged with a Nikon 90i Eclipse microscope as described above.

### Fluorescence recovery after photobleaching (FRAP)

FRAP was performed on a Nikon A1R laser-scanning confocal inverted microscope as described above for live cell imaging of 3T3 fibroblasts transduced with mEGFP-CBX2 and induced with 500 ng/mL of doxycycline. Images were acquired every 2 seconds for 90 seconds (45 frames). The first five frames were collected before the bleach pulse for baseline fluorescence. A circular region of interest (ROI) with a radius of 0.5-1 μm was selected for bleaching puncta with 100% laser power (488 nm). Fluorescent intensities and images analysis was done using Fiji software. FRAP curves were generated as previously described^22^ using three step normalization. First, the mean intensity of the bleach spot and the whole nucleus at each time point was normalized to the respective pre-bleach baseline intensity. Second, the relative bleach spot intensity was normalized to the relative nuclear intensity. Finally, the difference between the double-normalized FRAP intensity before and at the first frame after bleach pulse was calculated and normalized to 100%. FRAP recovery measurements were averaged over 15 replicates spanning multiple cells. Immobile fraction was estimated as percent fluorescence intensity unrecovered at last frame.

### Hexanediol treatments

Live cell imaging was performed as described above for 3T3 fibroblasts transduced with mEGFP-CBX2 and induced with 500 ng/mL of doxycycline. Images were acquired every 8 seconds for 600 seconds (75 frames). After 1 minute and as image acquisition was ongoing, 1,6-hexanediol diluted in media was added to a final concentration of 10%. In control experiments, an equal volume of media alone is added. The time of 1,6-hexanediol addition is time 0” and the first frame after addition is time 16”. Image analysis was done using Fiji.

### Preparation of Cy5-labeled ligands and incorporation into condensates

For visualization of DNA incorporation into condensates, the G5E4 nucleosome-positioning array^23^ was labeled with Cy5. The G5E4 array was excised from pG5E4 by restriction digest with Asp718, ClaI, DdeI and DraIII and purified by PEG precipitation. The excised fragment was end labeled using Klenow Fragment (New England Biolabs) to incorporate Cy5-dCTP into the G5E4 array.

For visualization of RNA incorporation into condensates, templates for *in vitro* transcription of CAT7 RNA^25^ were generated. The DNA sequence encoding CAT7 was amplified from human genomic DNA using primers incorporating a T7 promoter and subsequently cloned into pUC19. DNA templates for *in vitro* transcription were prepared by SmaI digest of the pUC19 vector containing T7-CAT7, followed by ethanol precipitation. *In vitro* transcription was performed with the MEGAscript T7 kit (Ambion), incorporating trace Cy5-UTP into the reaction. *In vitro* transcription proceeded for 4 hours at 37°C, followed by digestion of template DNA with DNase I for 30 minutes at 37°C. RNA was purified using a MEGAclear kit (Ambion).

For visualization of polynucleosome and MLA polynucleosome incorporation into condensates, HeLa nucleosomes were isolated as previously described^37^ and MLA nucleosomes containing an H3K27me3 analog were assembled as described^24^. HeLa and MLA nucleosomes were assembled onto Cy5-labeled G5E4 nucleosome-positioning arrays by salt dialysis as previously described^38^. Proper assembly of polynucleosome arrays was confirmed by EcoRI digest to visualize mononucleosomes and HhaI digest to assess the extent of occupancy of the central core of the array lacking nucleosome positioning sequences.

To assess incorporation of Cy5-labeled ligands into *in vitro* protein condensates, purified mEGFP fusion proteins were diluted into buffer as described above. Cy5-labeled ligands were added to pre-formed condensates to a final concentration of 0.3 μM. Contemporaneous incorporation of ligands into condensates was assessed by adding Cy5-labeled ligands to purified mEGFP fusion proteins prior to condensate formation. *In vitro* condensates were visualized by fluorescence microscopy as described above.

### Analysis of protein disorder and charge

Predicted protein disorder for CBX2 was calculated using the PONDR VSL2 algorithm^39^. Protein charge distribution was calculated for CBX2 variants using the EMBOSS charge algorithm^40^ with default parameters using a window size of 10 residues.

### Immunoblot analysis

The indicated cell lines were induced with the specified concentration of doxycycline for 24 hours, cells were lysed in RIPA buffer (Thermo Fisher Scientific, 89900), and protein was quantified by Bradford assay. Samples were run on SDS 4-20% polyacrylamide gels (Biorad) and transferred to nitrocellulose membranes. After transfer, membranes were blocked with 5% milk in TBS with 0.1% Tween-20 for 1 hour at room temperature. Membranes were incubated with anti-CBX2 (Santa Cruz, sc19297, 1:500) or anti-GAPDH (Santa Cruz, sc32233, 1:2500) diluted in 2% milk in TBS with 0.1% Tween-20 overnight at 4°C. After washing three times with TBS with 0.1% Tween-20 for 5 minutes at room temperature, membranes were incubated with secondary antibody conjugated to HRP (1:20,000) diluted in 1% milk in TBS with 0.1% Tween-20 for 1 hour at room temperature. Membranes were washed three times with TBS with 0.1% Tween-20 for 5 minutes at room temperature and developed with SuperSignal West Pico PLUS Chemiluminescent Subtrate (Thermo Fisher Scientific, 34577) and imaged using a Chemidoc (Biorad) or film. Quantification was done using Fiji software and relative expression level was normalized to 1 for CBX2 at each doxycycline concentration. For analysis of proteins obtained by co-IP, membrane was incubated with anti-GFP-HRP (Abcam, ab184207, 1:10,000), anti-CBX2 (Santa Cruz, sc19297, 1:500), anti-RING1b (Bethyl, A302-869A, 1:5,000), or anti-PHC1 (Active Motif, 39723, 1:1,000) and processed as above (note the secondary antibody step was omitted for anti-GFP-HRP blotted membranes). All co-IP membranes were imaged using a Chemidoc (Biorad). To compare expression of different CBX paralogs, CJ7 mESCs and E11.5 *Cbx2*^+/+^, *Cbx2*^+/-^, and *Cbx2*^−/−^ mouse embryos were examined in comparison to 3T3 fibroblasts. Embryos were homogenized by running through a 25G needle >10 times using a syringe and lysates were generated as above. Membranes were first stained with Ponceau prior to incubation with anti-CBX2 (Santa Cruz, sc19297, 1:500), anti-CBX4 (Millipore, MAB11012, 1:2000), anti-CBX7 (Santa Cruz, sc376274, 1:1000), or anti-CBX8 (Bethyl, A300-882A, 1:3000) and processed as above and imaged on film.

## Data Availability

The data that support the findings of this study are available from the corresponding author upon reasonable request.

## End notes

**Supplementary Information** is available in the online version of the paper.

## Acknowledgments

We thank G. Narlikar for advice initiating this project; R. Tomaino at the Taplin Mass Spectrometry Facility for all mass spectrometry analysis; J. Lee laboratory, the MGH Program in Membrane Biology (PMB) microscopy core, and M. Alimova at the Broad Institute of Harvard and MIT for help with and access to microscopes; L. Rubin laboratory for imaging software; the R.E.K. laboratory for fruitful discussions; and M.B. Ardehali, E. Jaensch, T. Oei, I. Tchasovnikarova and C. Tsokos for critical reading of the manuscript. This work was supported by the NIH (F32GM109693 to A.J.P. and R01GM04390 to R.E.K.) and graduate fellowships from the National Science Foundation and Albert J. Ryan Foundation to C.P.D.

## Author contributions

A.J.P., C.P.D. and R.E.K. designed the project. A.J.P. and C.P.D. created plasmids, purified proteins and did all sample preparation for microscopy and mass spectrometry. A.J.P. conducted all *in vitro* work, microscopy and Co-IPs. C.P.D. created cell lines and performed protein domain analysis. A.J.P. and J.K. performed immunoblotting. J.K. performed immunofluorescence. G.R. performed high-content imaging analysis. M.M.K. provided the CKII plasmid and advised on phosphorylation experiments. S.K.M generated MLA nucleosomes and labeled DNA constructs for arrays. A.J.P., C.P.D. and R.E.K. wrote the manuscript.

## Author Information

The authors declare no competing interests. Correspondence and requests for materials should be addressed to R.E.K.

**Extended Data Figure 1:**
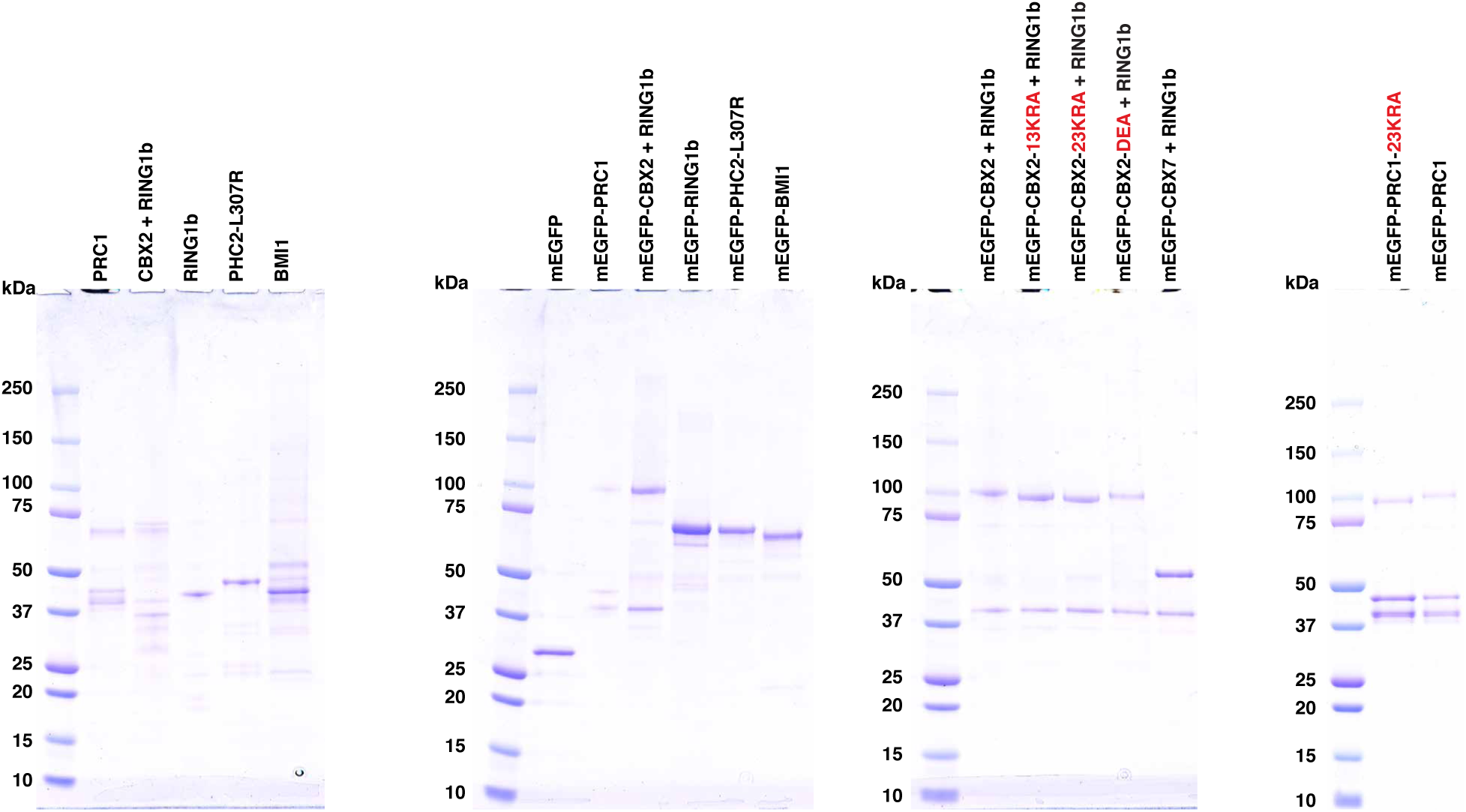
Purified PRC1 proteins. Panels from left to right are Coomassie stained SDS-polyacrylamide gels of purified recombinant untagged PRC1 subunits, purified recombinant mEGFP-tagged PRC1 subunits, purified recombinant mEGFP-tagged CBX + RING1b heterodimers and purified recombinant mEGFP-tagged PRC1 complexes.

**Extended Data Figure 2:**
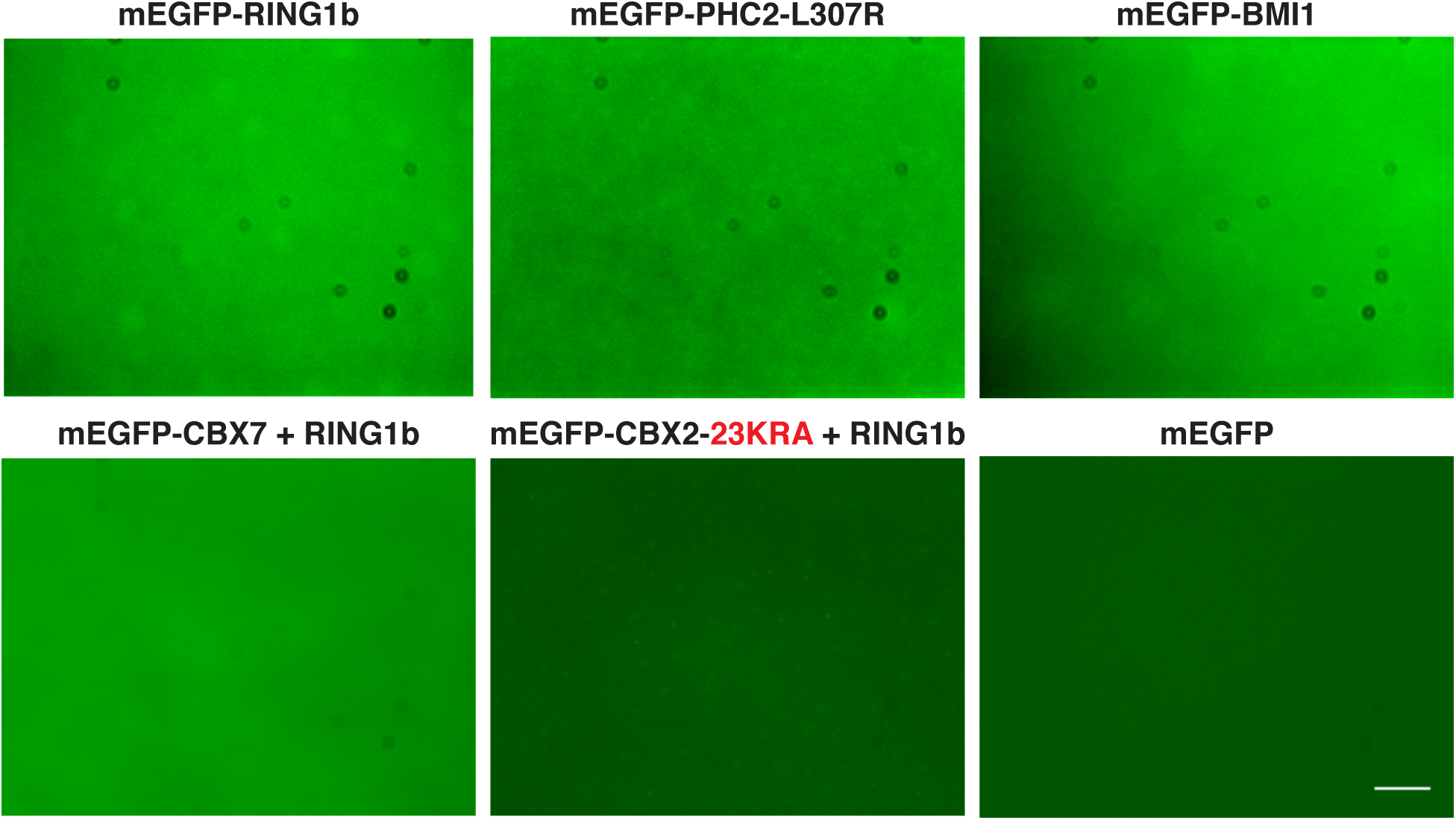
Higher concentrations of individual PRC1 subunits that do not phase separate *in vitro*. Micrographs of mEGFP-tagged PRC1 subunits in buffer containing 20 mM HEPES pH 7.9, 100 mM KCl, 1 mM MgSO_4_ at the following protein concentrations: RING1b, BMI1, PHC2-L307R, and mEGFP (12.5 μM), CBX2-23KRA + RING1b (18.4 μM), and CBX7 + RING1b (30 μM). For each experiment a representative micrograph from two independent protein preparations is shown. Scale bar = 10 μm.

**Extended Data Figure 3:**
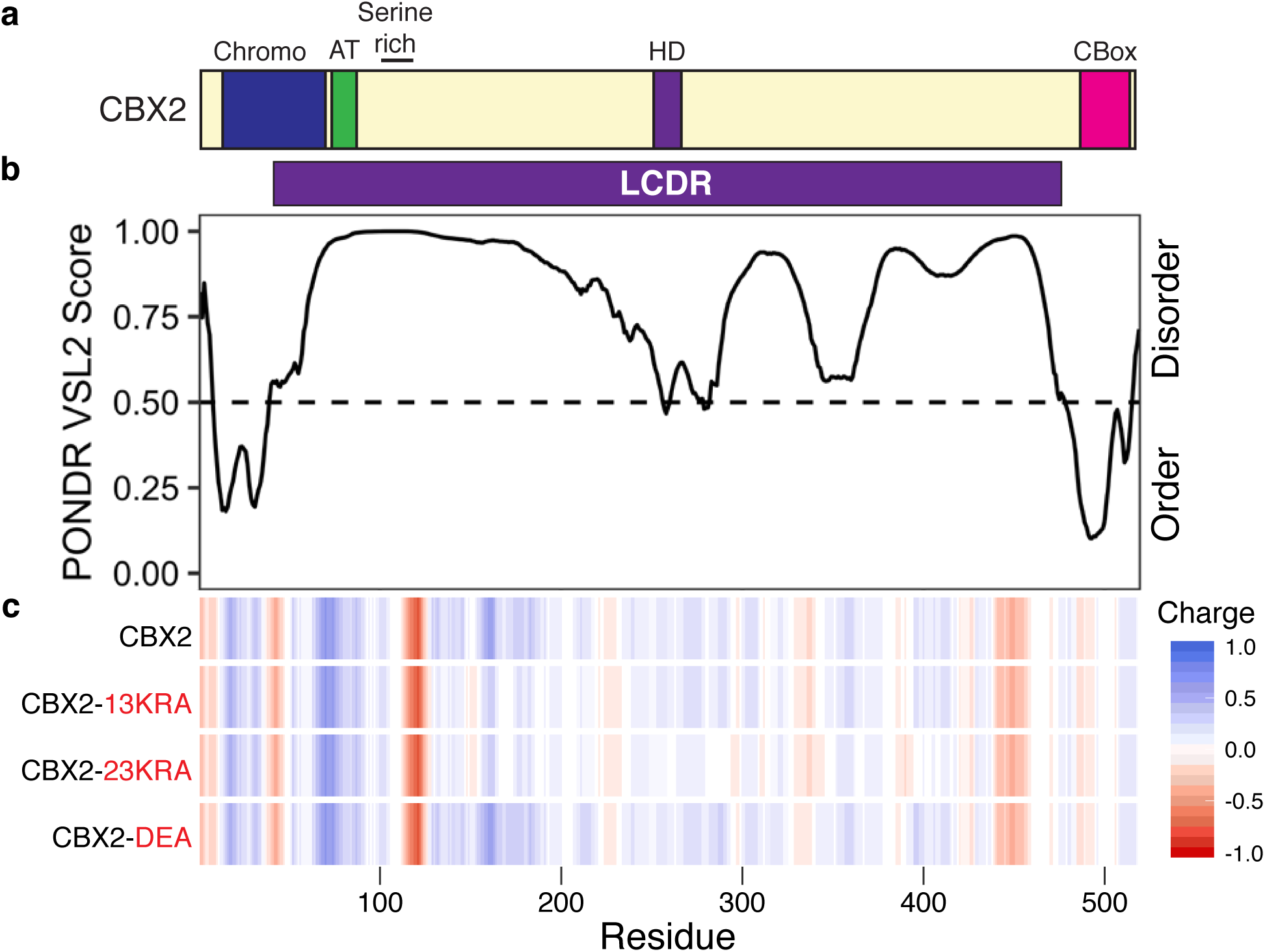
CBX2 contains a positively charged LCDR. **a**, Schematic of CBX2 protein domains. **b**, Graph plotting intrinsic disorder with Predictor of Natural Disordered Regions (PONDR) using the VSL2 algorithm for CBX2. Purple bar designates the LCDR in CBX2. **c**, Heat map indicating charge distribution across CBX2 for wild type and indicated charge mutants.

**Extended Data Figure 4:**
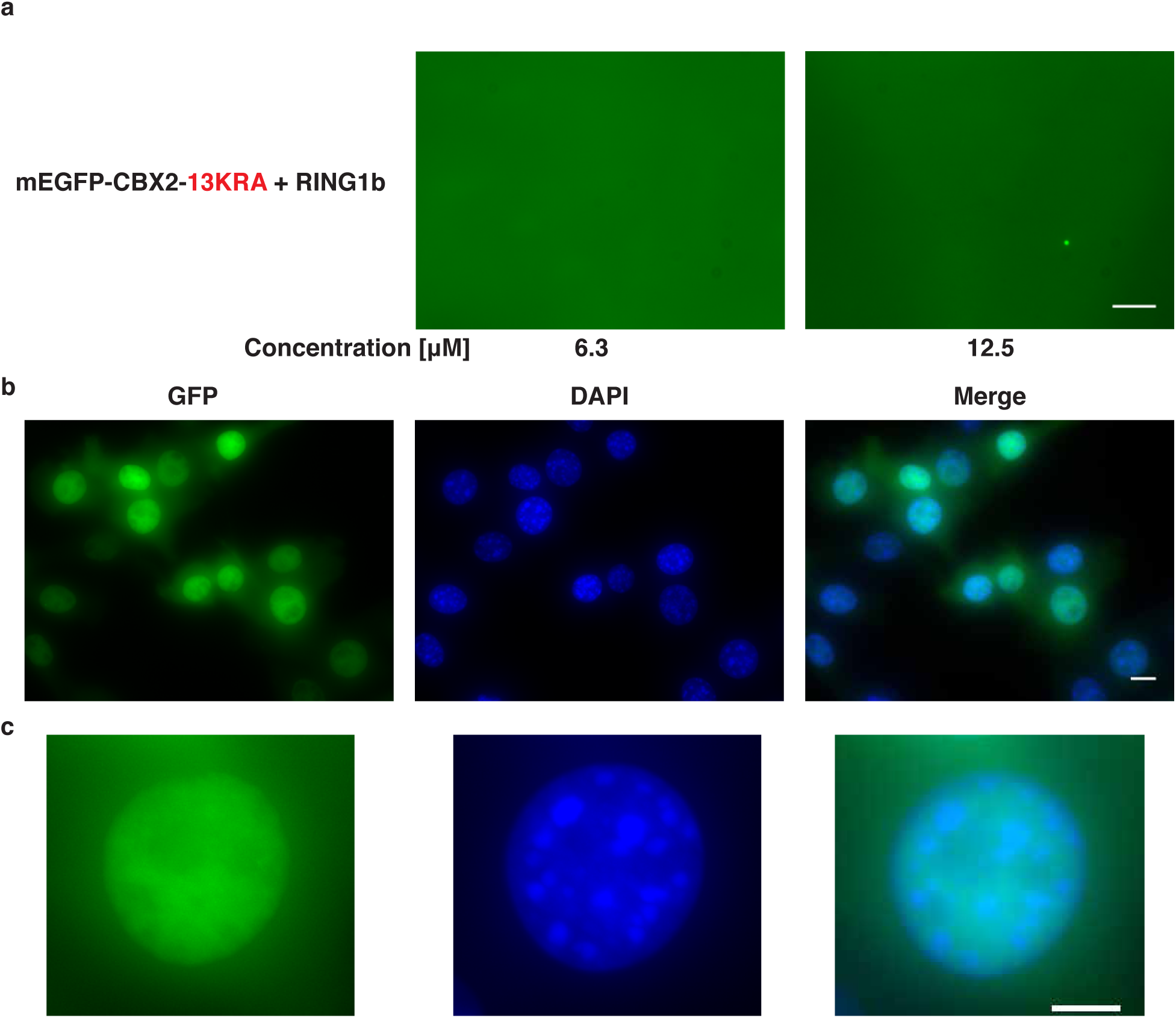
CBX2-13KRA does not phase separate in vitro and in vivo. **a**, Micrographs of mEGFP-CBX2-13KRA + RING1b at indicated protein concentration in buffer containing 20 mM HEPES pH 7.9, 100 mM KCl, 1 mM MgSO_4_. Scale bar = 10 μm. **b**, Representative micrographs of 3T3 fibroblasts expressing mEGFP-CBX2-13KRA after 500 ng/mL doxycycline induction. Column panels from left to right are the GFP channel, DAPI channel and merged images. Scale bar = 10 μm. **c**, Magnified images of representative nuclei from (b). All cell images are of Z-stacks using formaldehyde fixed cells. Scale bar = 5 μm.

**Extended Data Figure 5:**
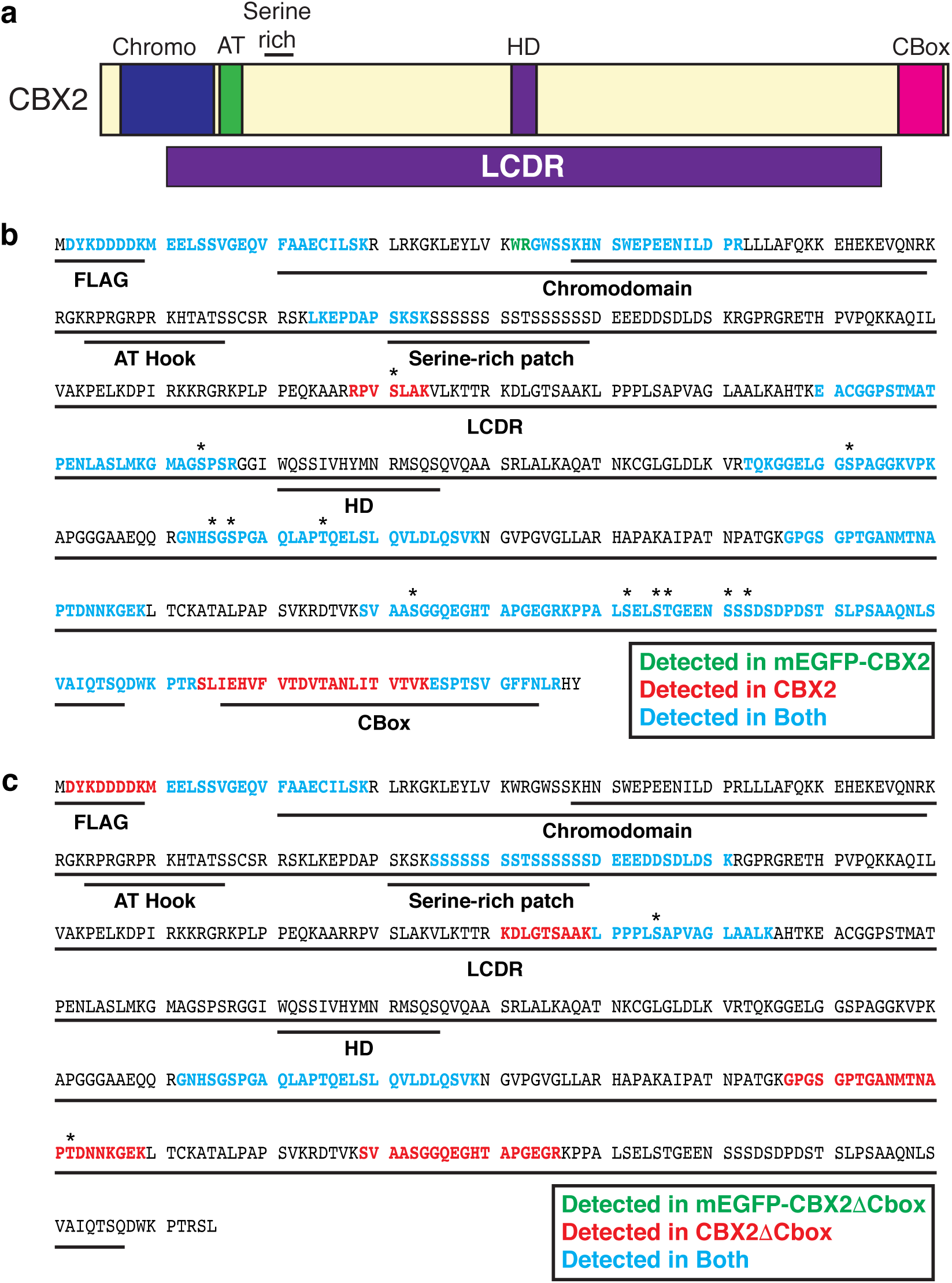
Phosphorylation sites in purified recombinant CBX2. **a**, Schematic of CBX2 protein domains with LCDR indicated. **b**, Schematic showing phospho-peptides and individual phosphorylated residues in CBX2 and mEGFP-CBX2 purified from Sf9 cells. Asterisks indicate residues that could be confidently called as phosphorylated within peptides containing multiple potential phosphorylation targets. **c**, Schematic showing phospho-peptides and individual phosphorylated residues in CBX2DCbox and mEGFP-CBX2DCbox purified from *E.coli* expressing CK2.

**Extended Data Figure 6:**
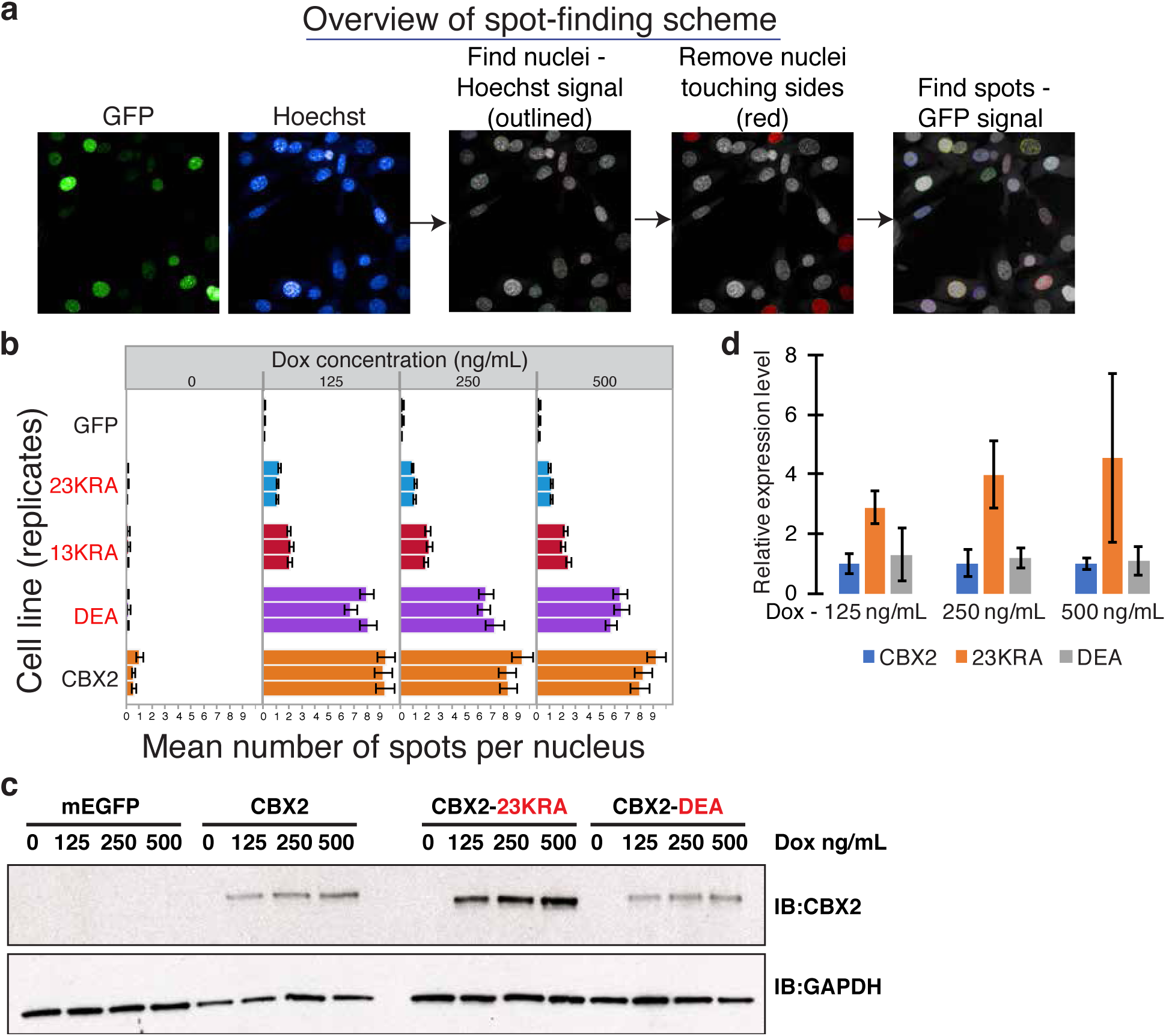
Punctate structure number and distribution are disrupted by CBX2-23KRA expression. **a**, Overview of spot-finding scheme used for (b) and (c). **b**, Quantification of mean number of spots per nucleus from three replicates for indicated 3T3 cell line expressing mEGFP-tagged CBX2 variants at indicated doxycycline concentration. **c**, Representative immunoblot showing expression level of indicated mEGFP-CBX2 variants in 3T3 fibroblasts at indicated doxycycline concentrations. GAPDH is shown for loading control. **d**, Quantification of relative protein expression level in two replicates of (c). Error bars represent standard deviation.

**Extended Data Figure 7:**
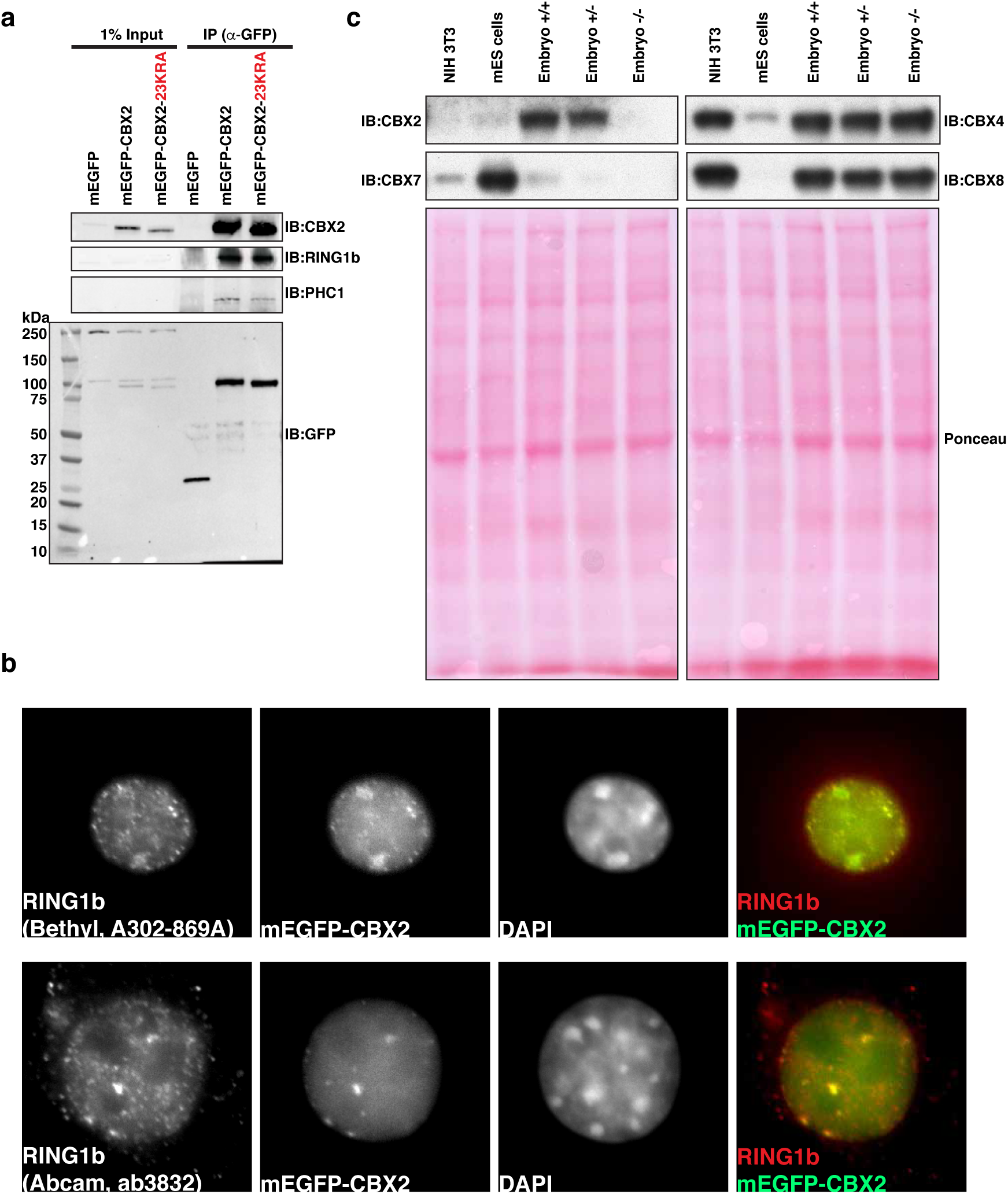
Induced mEGFP-CBX2 incorporates into PRC1 complexes. **a**, Co-immunoprecipitation of indicated mEGFP-tagged constructs and endogenous PRC1 subunits in 3T3 fibroblasts after 500 ng/mL doxycycline induction using anti-GFP antisera. **b**, Co-immunofluorescence of RING1b using indicated commercially available antibodies in 3T3 fibroblasts after 500 ng/mL doxycycline induction of mEGFP-CBX2. Column panels from left to right are the Cy3 channel (RING1b), GFP channel, DAPI channel and merged images of Cy3 and GFP. **c**, Immunoblot of indicated CBX homolog expression in NIH 3T3 fibroblasts, CJ7 mouse embryonic stem cells, WT mouse embryos, Cbx2 heterozygous and homozygous mutant mouse embryos. Ponceau staining of blots are shown for loading.

**Extended Data Figure 8:**
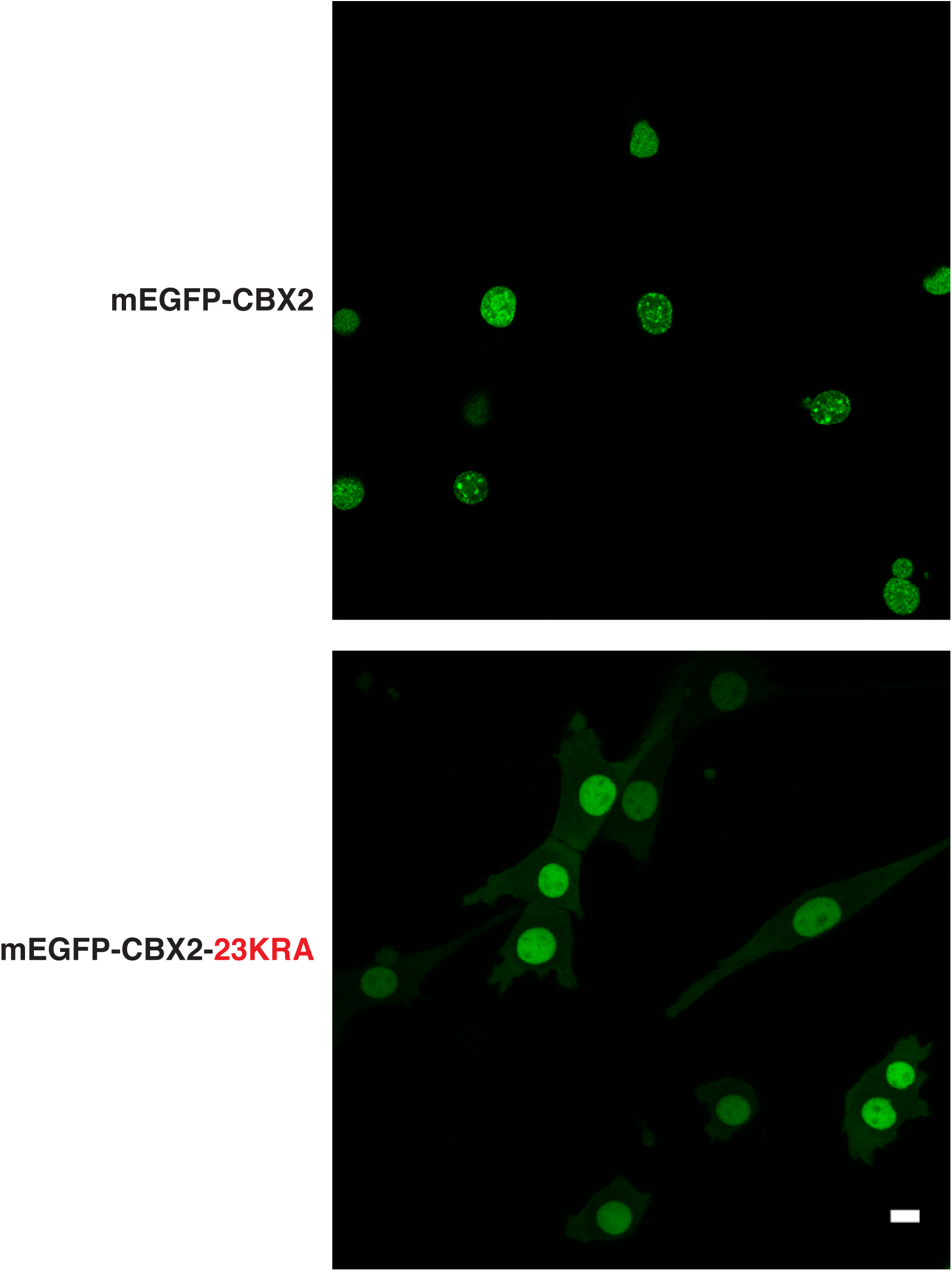
PRC1 forms punctate structures in live cells. Representative micrographs of live 3T3 fibroblasts expressing mEGFP-CBX2 or mEGFP-CBX2-23KRA after 500 ng/mL doxycycline induction. Scale bar = 10 μm.

**Extended Data Figure 9:**
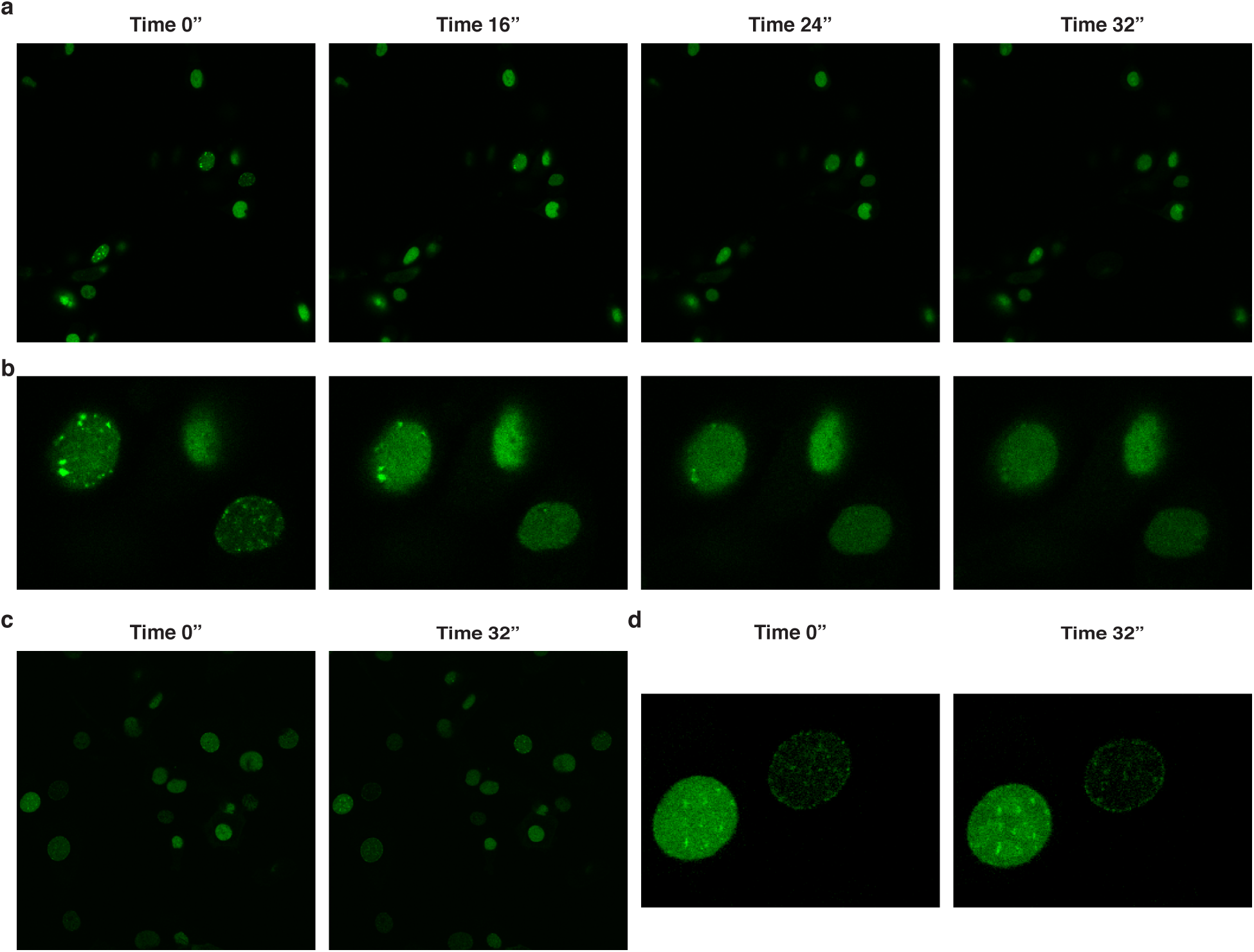
mEGFP-CBX2 puncta are disrupted by 1,6-hexanediol. **a**, Representative images of 1,6-hexanediol experiment at indicated times with mEGFP-CBX2 after 500 ng/mL doxycycline induction in 3T3 fibroblasts. Time 0” indicates when 1,6-hexanediol is added. **b**, Magnified images of representative nuclei from (a) showing loss of puncta upon 1,6-hexanediol addition. **c**, Control experiment with media lacking 1,6-hexanediol added and imaged at indicated times. **d**, Magnified images of representative nuclei from (c) showing retention of puncta.

**Extended Data Figure 10:**
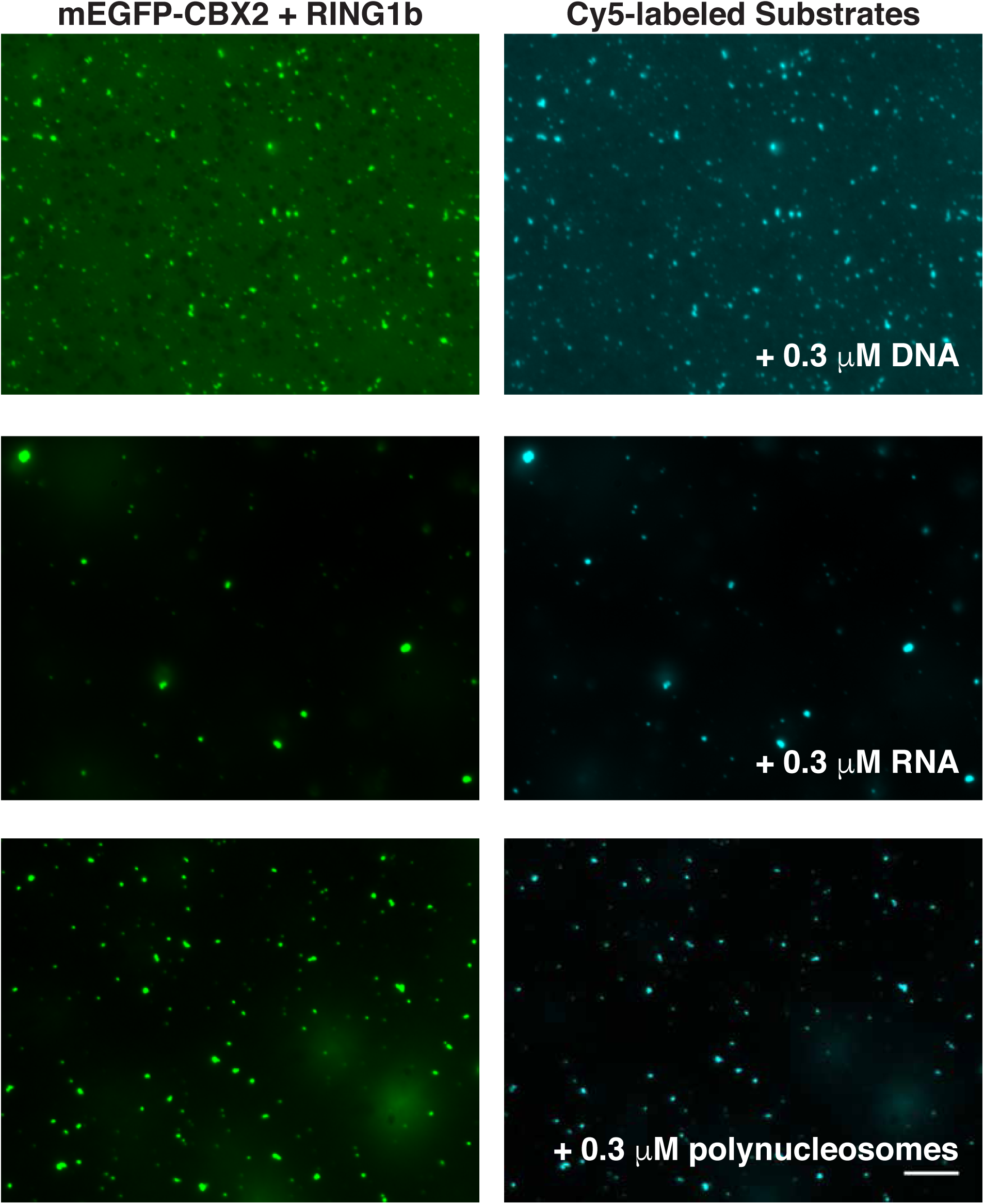
PRC1 ligands partition into condensates with mEGFP-CBX2 + RING1b when added after droplet formation. Micrographs of mEGFP-CBX2 + RING1b (6.3 μM) with indicated Cy5-labeled substrate (0.3 μM). Left panels are GFP channel and right panels are Cy5 channel. Cy5-labeled substrates were added after formation of mEGFP-CBX2 + RING1b droplets.

---

**Supplementary Table 1. (separate Excel file).** Phosphorylated peptides identified by mass spectrometry of purified CBX2.

**Supplementary Table 2. (separate Excel file).** PRC1 peptides recovered after co-IP mass spectrometry.

